# *metID:* A R package for automatable compound annotation for LC−MS-based data

**DOI:** 10.1101/2021.05.08.443258

**Authors:** Xiaotao Shen, Si Wu, Liang Liang, Songjie Chen, Kévin Contrepois, Zheng-Jiang Zhu, Michael Snyder

**Affiliations:** Department of Genetics, Stanford University School of Medicine, Stanford, CA, 94304, USA; Interdisciplinary Research Center on Biology and Chemistry, Shanghai Institute of Organic Chemistry, Chinese Academy of Sciences, Shanghai 200032, China

**Author notes:** These authors contributed equally (Xiaotao Shen, Si Wu).

## Abstract

**Summary:** Accurate and efficient compound annotation is a long-standing challenge for LC−MS-based data (e.g. untargeted metabolomics and exposomics). Substantial efforts have been devoted to overcoming this obstacle, whereas current tools are limited by the sources of spectral information used (in-house and public databases) and are not automated and streamlined. Therefore, we developed *metID*, an R package that combines information from all major databases for comprehensive and streamlined compound annotation. *metID* is a flexible, simple, and powerful tool that can be installed on all platforms, allowing the compound annotation process to be fully automatic and reproducible. A detailed tutorial and a case study are provided in Supplementary Materials.

**Availability and implementation:** https://jaspershen.github.io/metID/.

## 1 Introduction

Liquid chromatography coupled to mass spectrometry (LC−MS) is a comprehensive, unbiased technology to research small compounds, which has become increasingly popular in food, environment, and biomedical studies (Wishart, 2016)(Fraga-Corral *et al*., 2020). This approach has been very successful however the generated data has typically not been used to its full potential in part due to an incomplete annotation of the chemicals detected. Implementing an accurate and efficient compound annotation workflow would be invaluable to assist biological hypothesis generation and data interpretation. In order to standardize data reporting from metabolomics studies, the community proposed to adopt a grading scheme for annotation confidence ranging from 1-4 (Sumner *et al*., 2007) using the following parameters: mass to charge ratio (*m/z*), retention time (RT), and MS^2^ spectral matching. Confident annotation being the Achilles’ heel of modern metabolomics experiments, recent initiatives are now focusing on generating high-quality MS^2^ spectral libraries that are publicly accessible. By taking advantage of these resources, some tools (Chaleckis *et al*., 2019) were developed to facilitate compound annotation. However, these tools are limited by: (1) the breadth of parameters used to compute an annotation score; previous tools typically only use *m/z* and/or MS^2^ spectra information but lack RT; (2) the inability to combine spectral information from multiple sources including in-house and public databases in a streamlined fashion; (3) the processing speed, previous tools normally do not allow the implementation on cluster servers. More detailed comparisons between the existed tools and *metID* are listed in **Table S1**.

In this context, we developed a new R package, *metID*, particularly designed to (1) streamline the construction of users’ in-house databases from authentic chemical standards run in each laboratory and (2) automated compound annotation pipeline. As annotation level 1, *metID* provides our spectral and RT data from our in-house database containing more than 1,000 authentic standards acquired in HILIC and RPLC modes (Contrepois et al., 2015) (users need to use the same LC gradient to match RT). For level 2, *metID* now provides 5 public MS^2^ databases and 5 MS^1^ databases for level 3 (**Table S2**).

## 2 Features and methods

Using *metID*, users can easily build their in-house databases using the data from authentic standards that were acquired in their own laboratories. The public databases can also be easily organized as the database format for *metID*. The in-house databases in our laboratory and several public databases are provided as reference. *metID* can be installed on all platforms (Mac OS, Windows, and Linux).

### 2.1 Database construction for *metID*

If users have in-house standards which have been acquired with RT and MS^2^ spectra, it is possible to build the in-house database using the “construct_database()” function (**Fig. 1a**). Three items are contained in the database: (1) database information, (2) standard information, and (3) MS^2^ spectrum for each compound.

**Fig. 1.**
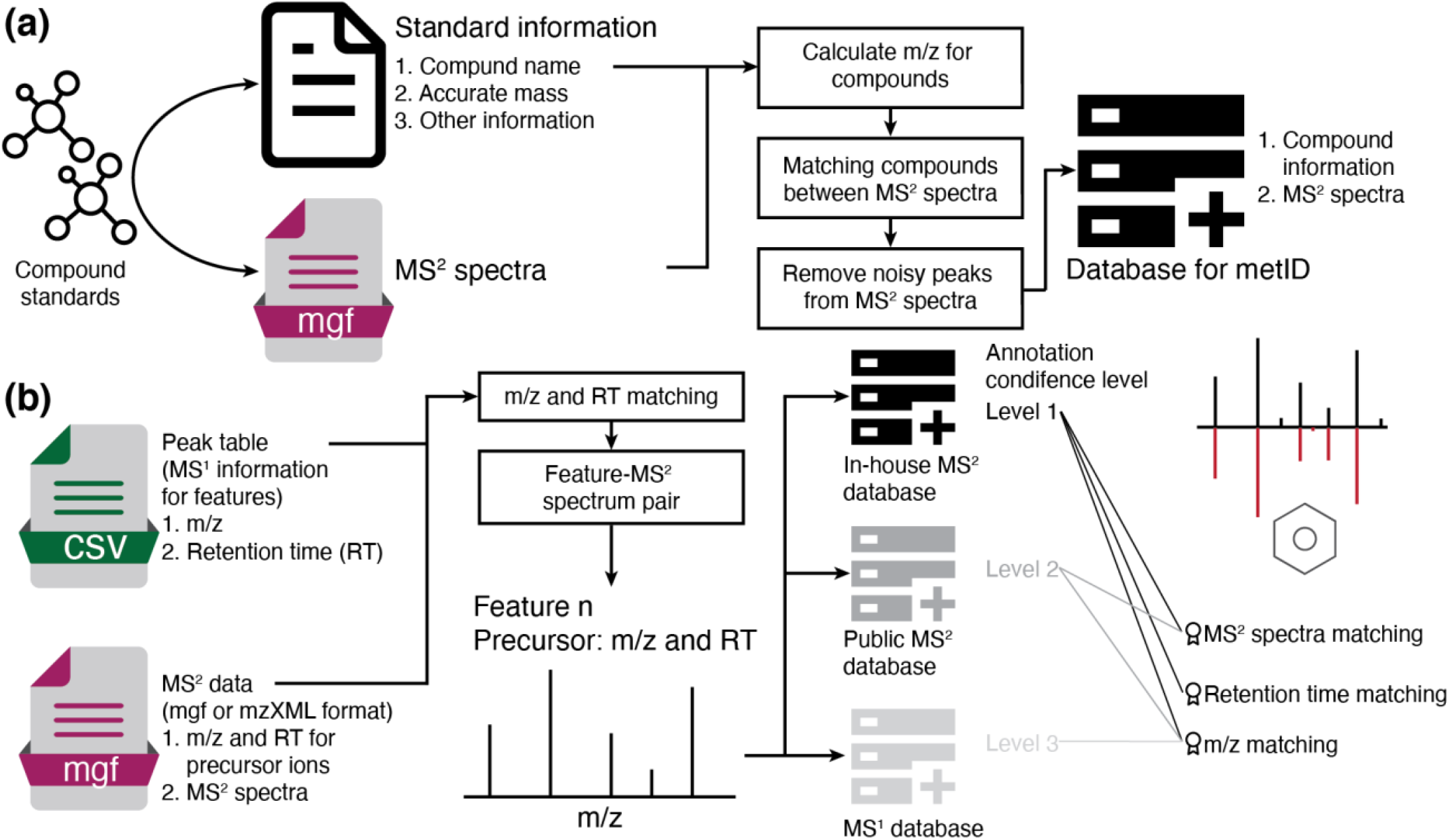
The overview of (a) in-house database construction and (b) compound annotation in different levels.

### 2.2 Compound annotation

RT may shift in different acquisition batches. Therefore, it is necessary to correct the RT in databases using the “correct_database_rt()” function if you spike internal standards into standards and subject samples. Then *metID* can annotate compounds with different levels according to databases (in-house database, level 1; public MS^2^ database, level 2; MS^1^ database, level 3). For *m/z*, RT, and MS^2^ spectra match scores, they are combined as one total score and scaled to 0-1 (Shen *et al*., 2019) (**Fig. 1b** and **Supplementary Material**).

## 3 Case study

We applied *metID* on a published study from our lab (Contrepois *et al*., 2020) as a case study to demonstrate the value of *metID* for automatic metabolite annotation (only ∼2 hour running on a Mac with 6-core 32 G memory). The authors reported 463 annotated metabolites with level 1-2 by manual inspection of *m/z*, RT, and MS^2^ (Level 1 as golden standards). By using *metID*, we successfully annotated 942 metabolites with high confidence (level 1-2). Comparing the annotation results from metID with the original annotation in the publication, *metID* retrieved all the 463 annotated metabolites in the previous paper in an automatic way. Besides these overlapping annotations, *metID* is able to annotate 479 new metabolites that were not annotated in the original publication (**Fig. S5**). These results indicate that *metID* is not only a valid approach for compound annotation with high accuracy (100%) and high speed (∼2 hours) but also a powerful tool to largely increase annotation coverage in an automatic way (**Supplementary Material**).

## 4 Conclusion

*metID* is used for compound annotation in untargeted LC−MS-based data (metabolomics and exposomics). It allows users to build and share their in-house databases and search against all the major spectral databases. To our best knowledge, it is the first R package to allow users to build in-house databases and combine them with the public databases for compound annotation and can be applied in all computer platforms automatically. As R is very popular in the bioinformatics field, it means that as an open-source tool, *metID* can contribute to all community members to increase its public databases and add new methods and steps to it.

## Funding

This work received no external funding.

## Conflict of Interest

M.S. is a co-founder and member of the scientific advisory boards of the following: Personalis, SensOmics, Filtricine, Qbio, January, Mirvie, and Oralome.

## Supplementary Information for

### 1. Comparison of existing annotation tools

The subjective assessment of the different annotation tools was made according to previous reports (Wen *et al*., 2017)(Chong *et al*., 2018) and our own experience (**Table S1**). The following criteria were used for the comparison:

1. “Incorporation of RT information” evaluates whether the tool allows incorporating retention time (RT) information from the in-house databases.
2. “Usage of MS^2^ spectrum data” is determined by whether the tool provides the metabolite annotation by using MS^2^ spectrum data.
3. “Enriched MS^2^ spectrum databases” evaluates how many MS^2^ spectrum databases the tool uses. If the tool uses more than three public MS^2^ databases, such as NIST, METLIN, MassBank, HMDB, etc, this aspect is labeled as “+++”. If the tool only uses a limited number of public libraries, for example, MAIT only uses HMDB, this aspect is labeled as “+”. If the tool does not use any MS^2^ spectrum databases, this aspect is labeled as “-”.
4. “Built package” is determined by whether this tool has been developed as an available public R or python package in a programming way.
5. “Automation” evaluates whether the tool is automated by a pipeline or is more only with the procedure. If the tool has a modularized pipeline that allows automated usage, the aspect is labeled as “√”. If the tool only has the concepts and description of the procedure but has not built the modularized pipeline, such as the approach by application of correlated ion information (Lynn *et al*., 2015), the aspect is labeled as “-”.
6. “Suit for large-scale metabolomics” assesses whether the tool has the capability to conduct the metabolite annotation for the large-scale metabolomics dataset. If the tool allows the running for the large-scale metabolomics with built R or Python package, the aspect is labeled as “√”; If not, it is labeled as “-”

**Table S1.** The main features manifested in *metID* compared to other existing tools.

| Tool name | *metID* | CliqueMS | MAIT | xMSannotator | Application of correlated ion information<sup>1</sup> |
| --- | --- | --- | --- | --- | --- |
| Incorporation of RT information | √ | - | - | - | - |
| Usage of MS<sup>2</sup> spectrum data | √ | - | √ | - | √ |
| Enriched MS<sup>2</sup> spectrum databases | +++ | - | + | - | +++ |
| Built in a package | √ | √ | √ | √ | - |
| Automated | √ | √ | √ | √ | - |
| Suitable for large-scale metabolomics | √ | √ | √ | √ | - |

Symbols used for feature evaluations with “√” for present, “-” for absent, and “+” for a more quantitative assessment, with more “+” indicating better support.

### 2. Supplementary Methods

*metID* is a highly flexible open-source tool written in R that combines in-house as well as public compound and MS^2^ spectral databases. Users are allowed to build and upload their own in-house databases including accurate mass, retention time, and MS^2^ spectra to achieve the highest confidence level 1. Public MS^2^ spectral data are used to achieve annotation confidence level 2, and compound databases only with accurate mass information serve as annotation confidence level 3. The demo in-house database and public databases can be accessed and downloaded at the *metID* website (https://jaspershen.github.io/metID/articles/public_databases.html), and customized data analysis steps can also easily be integrated. *metID* is available as an R package that can be downloaded from GitHub and used on all computer platforms.

#### 2.1 Database construction

The first level of annotation confidence requires comparing chemical standards with their accurate mass, retention time, and MS^2^ spectra (Sumner *et al*., 2007; Goodacre *et al*., 2007). If users have their in-house standards which have been acquired with MS^2^ spectra data, it is encouraged to construct their own in-house databases using the *metID* package.

##### 2.1.1 Data preparation

Users first need to transform the raw standard MS data (e.g., RAW data format for Thermo Fisher) from positive and negative modes to mzXML format by using chromatography conversion tools, such as Proteowizard MSConvert tool (http://proteowizard.sourceforge.net/) (Chambers *et al*., 2012). The parameter setting is shown in **Fig. S1**.

**Figure S1.**
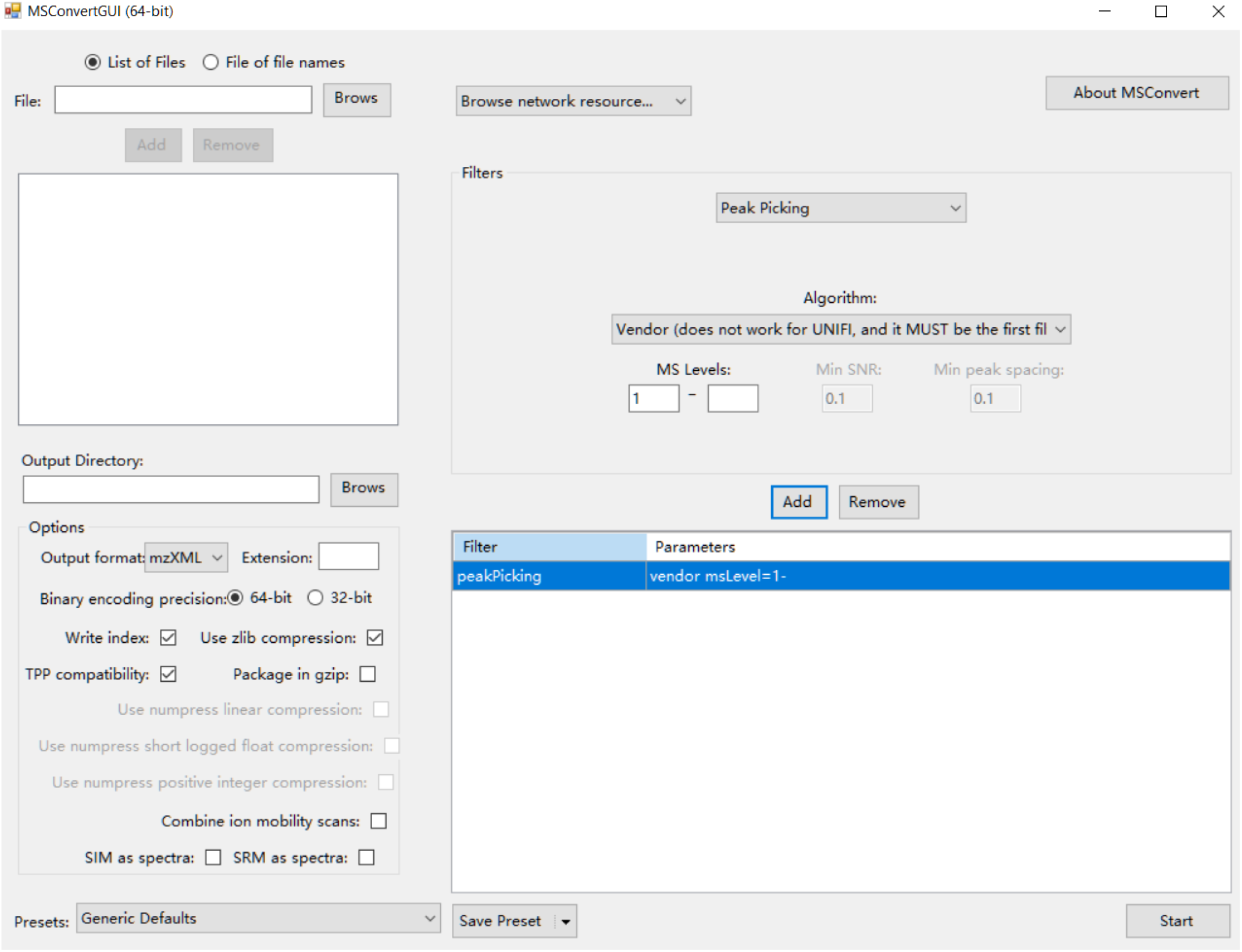
Parameter setting for MSConvert to convert raw MS data.

##### 2.1.2 Data organization

After that, *metID* requires users to organize compounds’ standard information in a table, and output with a CSV or xlsx format, which can refer to our demo data stored in “*demoData*” package (https://github.com/jaspershen/demoData).

From column 1 to 11, the columns are “Lab.ID”, “*mz*”, “RT”, “CAS.ID”, “HMDB.ID”, “KEGG.ID”, “Formula”, “mz.pos”, “mz.neg”, “Submitter”, respectively. It is allowed to have extra information for compounds in the in-house library, and users are able to compile them in the additional columns after column #11. Like the demo data shows, there is other additional information, namely “Family”, “Sub.pathway” and “Note”.

We encourage users to create a folder to hold mzXML format databases (e.g., positive and negative modes in “POS” and “NEG” folders, respectively) and the table with compound standard information.

##### 2.1.3 In-house database construction

We run the “construct_database()” function to build the in-house library for the first level of annotation searching. The arguments of “construct_database()” can be found using “?construct_database()”. We also provide the demo data in the “demoData” package to show how to construct the in-house library in *metID*.

As a demonstration, our lab collected and constructed an in-house library for 1,075 unique metabolites with reverse-phase liquid chromatography (RPLC) and hydrophilic interaction chromatography (HILIC). These 1,075 unique metabolites comprise different chemical classes, including amino acids (21.5%), carbohydrates (5.76%), lipids (38.89%), organic acids (9.39%), nucleotides (9.11%), cofactors and vitamins (3.44%), xenobiotics (8.09%), peptides (3.34%) and others (< 1%) (**Fig. S2**). To facilitate downstream analysis, such as pathway enrichment analysis and network analysis, we also curated these compounds with different database ID (e.g., HMDB, KEGG, CAS, etc), and 816 of them (76%) have the curated compound index from at least one public compound library (HMDB or KEGG), largely facilitating the application and reproducibility of metabolomics downstream analysis. The data from our in-house databases were acquired using Thermo Fisher QE-plus. However, the LC system may differ between labs. It is suggested to set rt.match.tol as 100000000 (no limitation) if users want to use our in-house library as demonstrated. The in-house database is provided in the *metID* website (https://jaspershen.github.io/metID/articles/public_databases.html).

**Figure S2.**
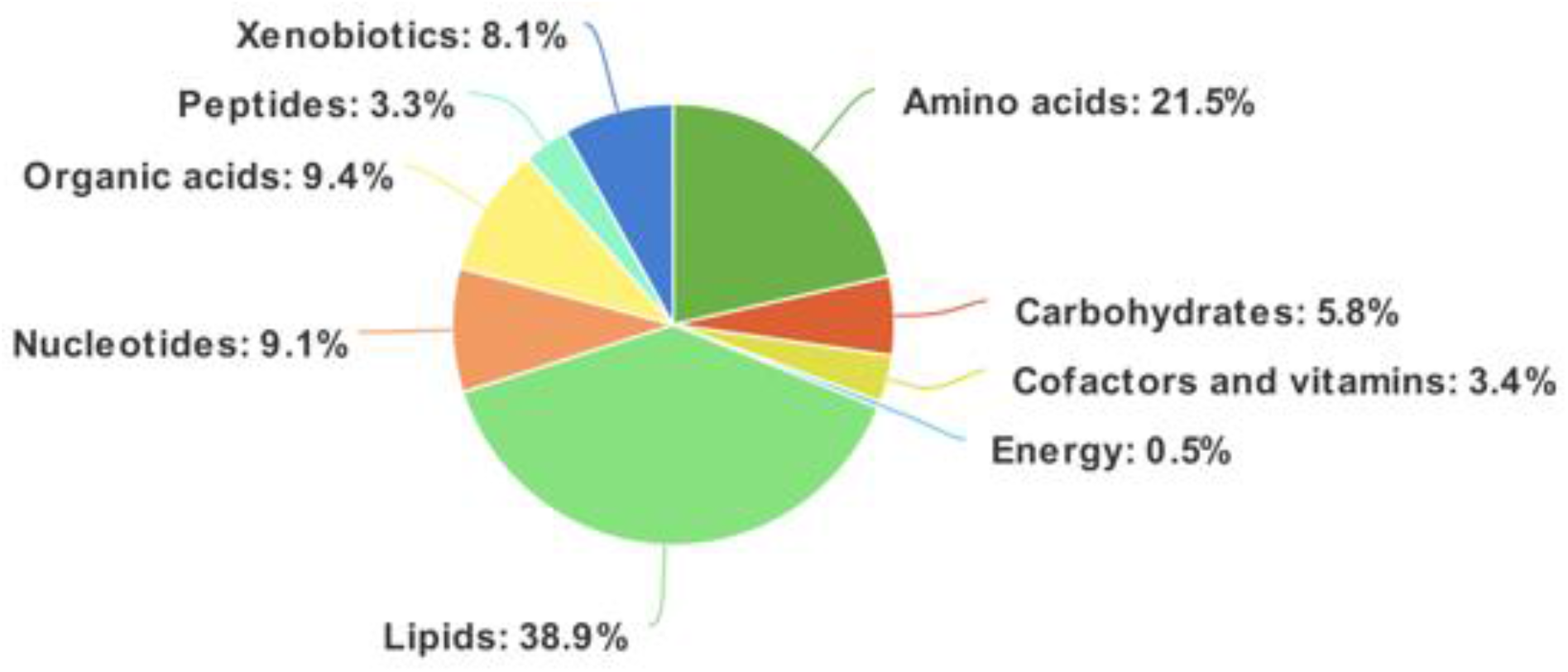
The chemical class of metabolites in the in-house library.

#### 2.2 Retention time correction

Metabolite retention time (RT) may shift in different acquisition batches. Therefore, it is necessary to correct the RTs in databases using the “construct_database()” function in *metID* before running the searching step, if you spike internal standards between the biological samples in the experiment.

First, users are required to prepare two internal standard (IS) tables for the in-house database and the actual biological samples with IS index, IS name, and their RT in columns. These two IS tables should be named separately, such as “database.is.table.xlsx” and “experiment.is.table.xlsx”. Then users can run the “construct_database()” function for RT correction of the constructed in-house database for every independent experiment. The main input for this function are two IS tables and the in-house database which you want to correct RTs. The output database with corrected RTs should be used for the following “metabolite identification” step.

#### 2.3 Metabolite identification

##### 2.3.1 Identify metabolites based on MS^1^ and MS^2^ databases

Users first need to prepare the peak table with MS^1^ data containing “name” (peak name), “mz” (mass to charge ratio), and “rt” (retention time, unit as second). It can be from any data processing software (such as XCMS (Smith *et al*., 2006), MS-DIAL (Tsugawa *et al*., 2015), etc). Next, the raw MS^2^ data from DDA (data-dependent acquisition) or DIA (data-independent acquisition) should be transformed to msp, mgf, or mzXML formats by using ProteoWizard MSConvert software (Chambers *et al*., 2012). Then users need to place the MS^1^ peak table, MS^2^ data, as well as the public MS^2^ databases which they want to use in one separate folder. The public MS^2^ databases provided in *metID* include HMDB (Wishart *et al*., 2018), MassBank (Horai *et al*., 2010), MoNA (Kind *et al*., 2018), Fiehn HILIC database (https://fiehnlab.ucdavis.edu/staff/kind/publications), and MassBank-orbitrap (Kind *et al*., 2018), DrugBankMS (Wishart *et al*., 2008), T3DB (Wishart *et al*., 2015), bloodExposomeMS1Database (Barupal and Fiehn, 2019) (**Table S2**).

After preparing and placing all these files in one folder, one can use the “*identify_metabolites*()” function to search against the in-house library that users create or upload and the public MS^2^ spectra databases that users select in *metID*. The arguments of “*identify_metabolites*” can be found using “?*identify_metabolites*”. *metID* allows users to choose the proper instrumental parameters according to their own instrument settings in the lab, for example, polarity, collision energy, column, adduct table, etc. Of note, one of the arguments called “ms2.match.tol” means the MS^2^ similarity tolerance for peak and database metabolite match. The MS^2^ similarity scores are calculated according to the algorithm from MS-DIAL (Tsugawa *et al*., 2015). This scoring metric and the “total score” can be used for further filtering steps and selection of best annotation for downstream analysis.

In brief, *metID* utilizes three scores to assess the matching between experimental peaks and compounds in databases, (1) accurate mass matching (m/z matching), (2) retention time (RT matching), and (2) MS^2^ spectra matching. For accurate mass matching and RT matching, the m/z error and RT error are calculated according to our previous study (Shen *et al*., 2019), and then convert to match scores with the linear transformation. The MS^2^ spectra matching algorithm is (cosine score) from MS-DIAL (Tsugawa *et al*., 2015) with little modification. Finally, all three scores are combined with different weights (default, m/z matching score: 0.25, RT matching score: 0.25, and MS^2^ spectra matching score: 0.5) as one total score.

The main output is a “*metIdentifyClass*” object, and users can print it out to see the identification information. It is also possible for users to set their own processing parameters by applying the “?” function, and check the corresponding processing parameters by using the “*get_parameters()*” function. Users can output the complete searching results without filtering in the identification table by applying the “*get_identification_table()*” function.

**Table S2.** The databases provided in *metID*.

| Database | Compound number | Spectrum number | Information<sup>1</sup> | Level<sup>2</sup> | Source |
| --- | --- | --- | --- | --- | --- |
| msDatabase_rplc0.0.2 | 917 | 2,047 | mz/RT/MS<sup>2</sup> | 1 | Snyde lab |
| msDatabase_hilic0.0.2 | 846 | 2,570 | mz/RT/MS<sup>2</sup> | 1 | Snyder lab |
| hmdbDatabase0.0.2 | 5,646 | 22,331 | mz/MS<sup>2</sup> | 2 | https://hmdb.ca/downloads |
| massbankDatabase0.0.2 | 18,311 | 91,148 | mz/MS<sup>2</sup> | 2 | https://massbank.eu/MassBank/ |
| monaDatabase0.0.2 | 19,537 | 40,315 | mz/MS<sup>2</sup> | 2 | https://mona.fiehnlab.ucdavis.edu/ |
| orbitrapDatabase0.0.1<sup>3</sup> | 8,360 | 23,227 | mz/MS<sup>2</sup> | 2 | https://mona.fiehnlab.ucdavis.edu/ |
| fiehn_hilic_database0.0.1 | 3,042 | 3,042 | mz/MS<sup>2</sup> | 2 | https://fiehnlab.ucdavis.edu/staff/kind/publications |
| hmdbMS1Database0.0.1 | 114,004 | 0 | mz | 3 | https://hmdb.ca/downloads |
| keggMS1Database_1.0 | 16,409 | 0 | mz | 3 | https://www.genome.jp/kegg/compound/ |
| drugbankMS1Database5.1.8 | 11,174 | 0 | mz | 3 | https://go.drugbank.com/releases/latest |
| T3DBMS1Database_1.0 | 3,533 | 0 | mz | 3 | http://www.t3db.ca/downloads |
| bloodExposomeMS1Database_1.0 | 65,957 | 0 | mz | 3 | https://bloodexposome.org/#/download |

1. The information of compounds in the database. 2. The annotation level used the databases for compound annotation. 3. This database includes all compounds that are acquired by Orbitrap in MoNA.

##### 2.3.2 Filtering

If users only want to review a subset of the identification results, it is possible to use the “*filter_identification*()” function to filter the complete results from the “*identify_metabolites*Class” object according to *m/z* error, rt error, MS^2^ spectra similarity, and total scores.

##### 2.3.3 MS^2^ spectra match plot output

Users can also use the “*ms2plot()*” function to visualize and output the MS^2^ spectra match plot for one, multiple, or all peaks. It is also possible to output interactive MS^2^ spectra match plots by setting “interaction.plot” as TRUE.

#### 2.4 Result integration and output

Users can choose any or all public MS^2^ spectra databases provided in *metID* to search against the peaks from their own experiment. We can use the “*identify_metabolites_params()*” function to edit the databases that we want to use for searching. Then we can run the “*identify_metabolites_all*()” function to conduct a searching process based on the databases that you select, it will return a list of objects of identification results.

Users can also annotate metabolites putatively only based on accurate mass, which is level 3 annotation according to MSI (Sumner *et al*., 2007; Goodacre *et al*., 2007). On the *metID* website, we provide 5 public MS^1^ databases (**Table S2**).

Of note, users can also use the “*identify_metabolites_all()*” function to run all the databases (in-house [level 1], public database [level 2] and MS^1^ database [level 3]) in one run.

### 3. Case study

To showcase the functionality of *metID*, we performed metabolite annotation using *metID* and compared the results to a published study where metabolite annotation was done mostly manually (Contrepois *et al*., 2020). Untargeted metabolomics data were generated on 310 human plasma using a combination of HILIC- and RPLC-MS approaches operated in both positive and negative ionization modes. The list of metabolites reported in the study are listed in **Supplementary Data1**. *metID* requires a peak table that includes a list of metabolic peaks with peak ID, annotated *m/z*, and retention time (RT). The generation of the peak table from LC-MS raw data can be achieved by using many software tools, such as XCMS, MZmine. According to the original publication for the metabolomics dataset as our case study, the raw data was processed by using Progenesis QI software (https://www.nonlinear.com/progenesis/qi/, v2.3, Nonlinear Dynamics). To ensure reproducibility, all the datasets (RPLC and HILIC in positive and negative modes) in the case study were provided.

#### 3.1 Case study information

The case study is a multi-omics study including a large-scale metabolomics experiment that aims to understand the molecular dynamics in response to acute exercise. More details about the metabolomics dataset in the case study are provided below.

**Table S3.** The information of the case study dataset.

| Variable | Information |
| --- | --- |
| Sample type | Plasma |
| Subject sample number | 310 |
| QC sample number | 36 |
| Data acquisition | Untargeted metabolomics |
| Instrument platform | LC-MS |
| Chromatographic condition | RPLC + HILIC |
| Polarity mode | Positive + negative |

#### 3.2 Sample preparation and LC-MS analysis

A detailed description of sample preparation, LC-MS analysis, and data processing was included in our previous publication (Contrepois *et al*., 2020). In brief, metabolites were extracted using 1:1:1 acetone:acetonitrile:methanol, evaporated to dryness under nitrogen, and reconstituted in 1:1 methanol:water before analysis. Metabolic extracts were analyzed four times using HILIC and RPLC separation in both positive and negative ionization modes. Data were acquired on a Q Exactive Plus mass spectrometer for HILIC and a Q Exactive mass spectrometer for RPLC (Thermo Scientific, San Jose, CA, USA). Both instruments were equipped with a HESI-II probe and operated in full MS scan mode. MS^2^ data were acquired on quality control samples (QC) consisting of an equimolar mixture of all samples in the study. HILIC experiments were performed using a ZIC-HILIC column 2.1 3 100 mm, 3.5 mm, 200A° (cat#1504470001, Millipore, Burlington, MA, USA) and mobile phase solvents consisting of 10 mM ammonium acetate in 50/50 acetonitrile/water (A) and 10 mM ammonium acetate in 95/5 acetonitrile/water (B). RPLC experiments were performed using a Zorbax SBaq column 2.1 3 50 mm, 1.7 mm, 100A° (cat# 827700-914, Agilent Technologies, Santa Clara, CA, USA) and mobile phase solvents consisting of 0.06% acetic acid in water (A) and 0.06% acetic acid in methanol (B). Data quality was ensured by (i) injecting 6 and 12 pooled samples to equilibrate the LC-MS system prior to run the sequence for RPLC and HILIC, respectively, (ii) injecting a pooled sample every ten injections to control for signal deviation with time, and (iii) checking mass accuracy, retention time and peak shape of internal standards in each sample.

#### 3.3 Data processing

Data from each mode were independently analyzed using Progenesis QI software (v2.3, Nonlinear Dynamics). Metabolic features from blanks and that did not show sufficient linearity upon dilution in QC samples (r <0.6) were discarded. Only metabolic features present in >2/3 of the samples were kept for further analysis. Inter- and intra-batch variation was corrected using the LOESS (locally estimated scatterplot smoothing local regression) normalization method on QC injected repetitively along with the batches (span = 0.75). Data were acquired in three and two batches for HILIC and RPLC modes, respectively. Missing values were imputed by drawing from a random distribution of low values in the corresponding sample (Tyanova et al., 2016).

#### 3.4 *metID* processing results

The preprocessed metabolomics data in the case study was utilized to demonstrate the performance of *metID*. All of the steps were conducted with R command lines according to the *metID* tutorial.

By comparing with the annotation results in the original publication (Contrepois *et al*., 2020), we retrieved all the 463 metabolites annotated in the original paper. However, we detected there are six metabolites that we got inconsistent annotation from the previous report (**Table S4**). With further investigation, we found that for one of these six metabolites with inconsistent annotation, metID also acquired the same annotation as that in the original paper, however, it was ranked as #2 based on the total score, while metID reported the top 1 annotation based on the default setting. The annotation of these top two metabolites pointed to the chemical isomers salicylic acid and hydroxybenzoic acid (**Figure S3)**, which are difficult to distinguish even with the MS^2^ spectrum and make the scores for these two metabolites very close. For the other five metabolites with the inconsistency of annotation, we found that the annotation by metID has higher scores than the annotation in the original paper, indicating metID has high annotation accuracy. The detailed Comparison of five inconsistent annotations between the exercise paper and metID is shown in **Figure S4**.

**Table S4.**
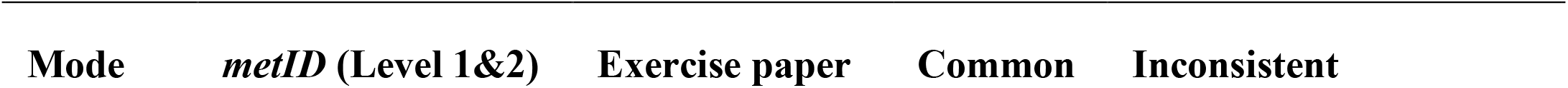
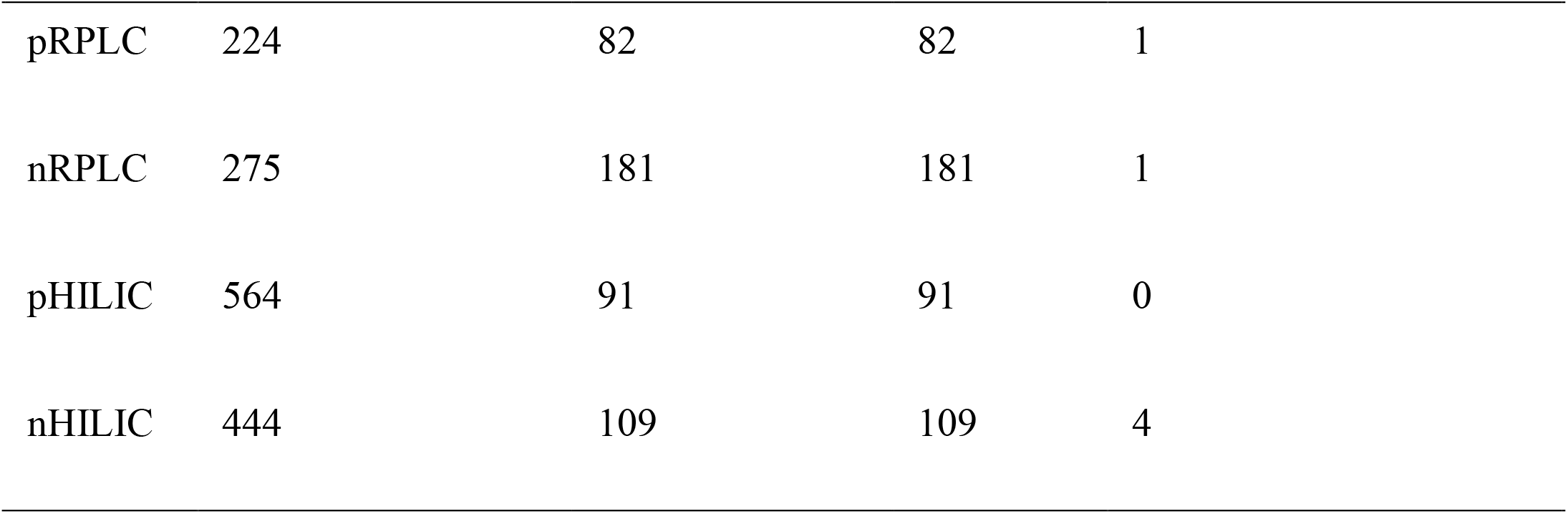
The comparison of annotation results by using *metID* and from the original publication.

**Figure S3.**
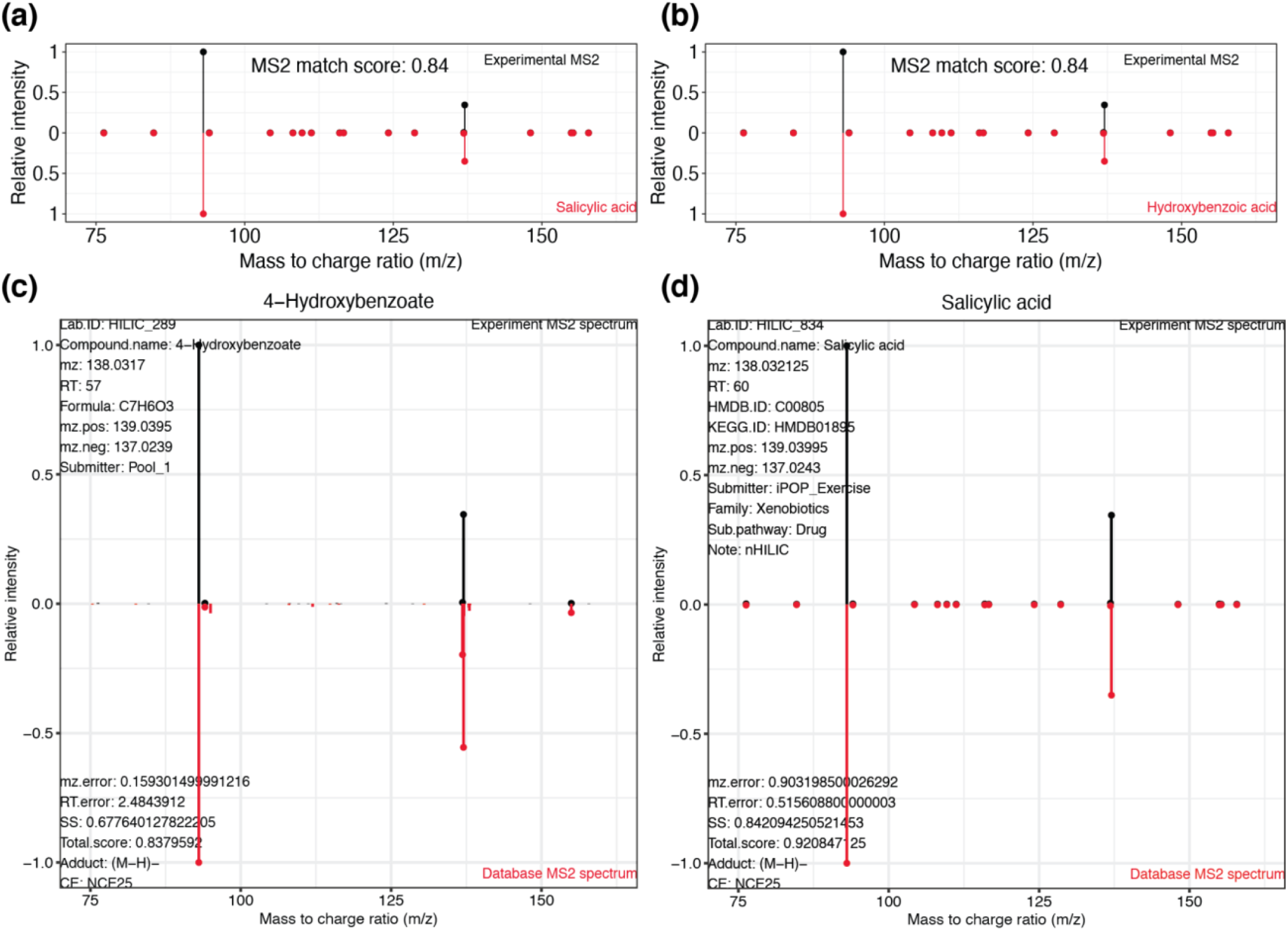
Annotation of peak 1.00_137.0243m/z with inconsistent annotation from the exercise paper. (a) The MS^2^ spectra matching plot between peak 1.00_137.0243m/z and standard salicylic acid (m/z error 0.90, retention time error 0.52 s, MS^2^ match score 0.84). (b) The MS^2^ spectra matching plot between peak 1.00_137.0243m/z and standard hydroxybenzoic acid (m/z error 0.90, retention time error 0.52 s, MS^2^ match score 0.84). The peak 1.00_137.0243m/z is also identified as 4-Hydroxybenzoate (rank 3, c) and Salicylic acid (rank 2, d). The top (black) MS^2^ spectrum is from the experiment and the bottom (red) MS^2^ spectrum is from the standard.

**Figure S4.**
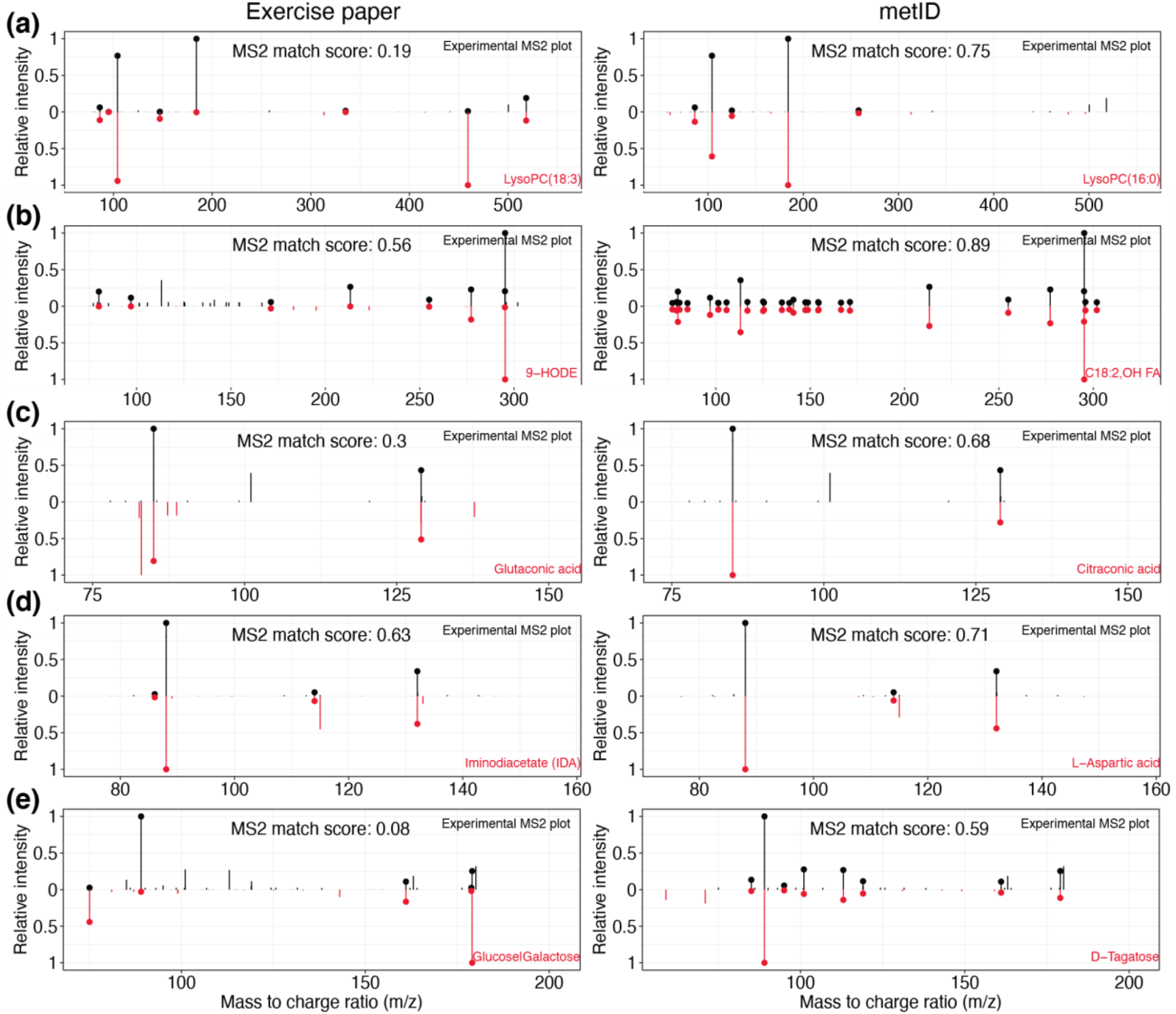
Comparison of five inconsistent annotations between the exercise paper and *metID*. (a) Peak 9.37_518.3231m/z, (b) peak 9.70_295.2272m/z, (c) peak 3.87_129.0193m/z, (d) 10.24_132.0302mz and (e) peak 7.88_180.0634n. Left columns are annotations from exercise paper and right columns are annotations from *metID*. Top (black) MS^2^ spectrum is from experiment and bottom (red) MS^2^ spectrum is from standard.

More importantly, we successfully annotated 479 unique additional metabolites by metID with level 1 and 2 that were not annotated in the original publication (Contrepois *et al*., 2020). The detailed comparison for each chromatographic condition in positive and negative modes was provided in **Supplementary Data 1**. With more investigation, the chemical composition for these 479 additional metabolites is shown in **Fig. S5**. In order to demonstrate the validity of the annotation for these additional metabolites, we presented the actual MS^2^ spectrum for these 479 newly annotated metabolites and the reference MS^2^ spectrum, as well as the comparison scores in

**Fig. S6.** These results show the advantages of *metID* in enlarging the annotation coverage with the guarantee of accuracy.

**Figure S5.**
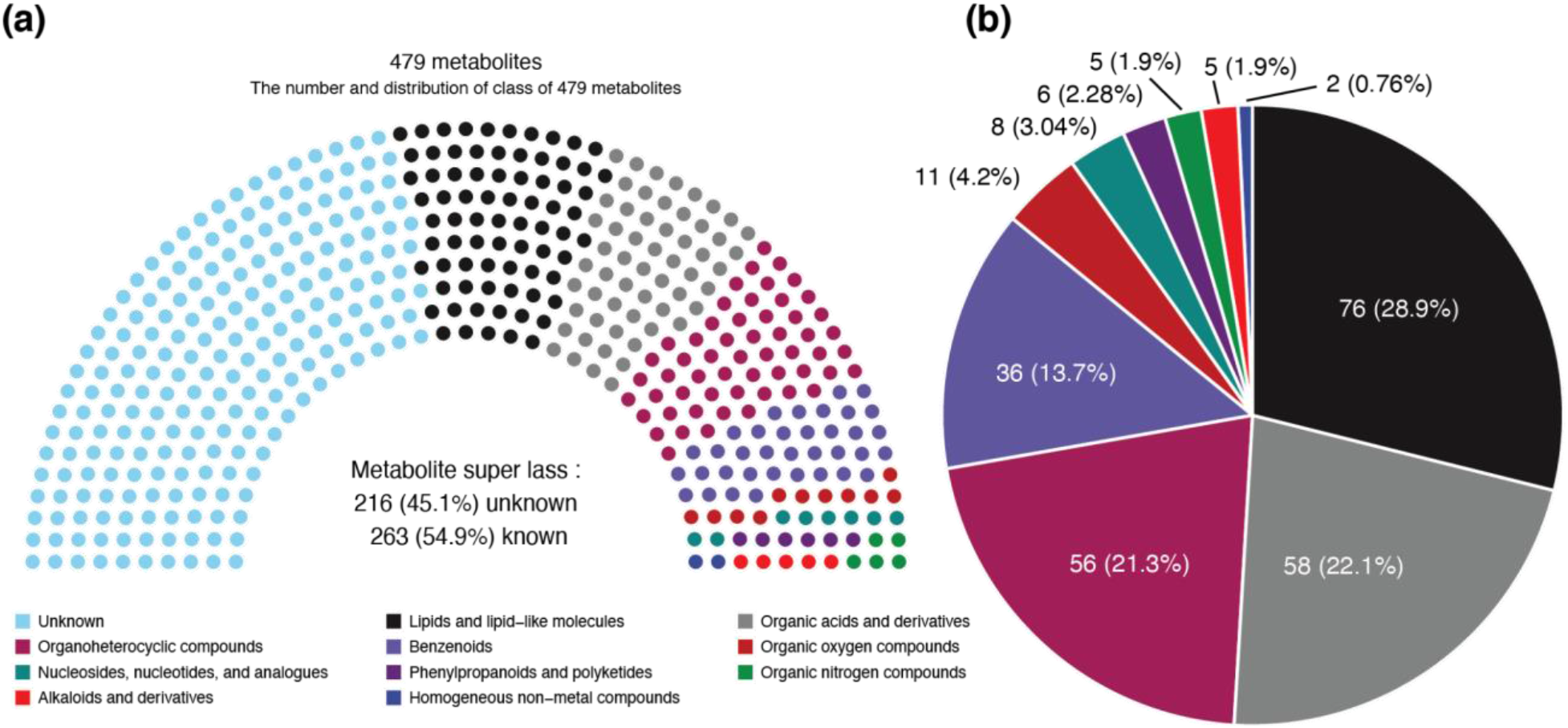
Super class of additional 479 metabolites from *metID*. (a) Classyfire was utilized to get the super class of metabolites. Among 479 metabolites, 216 (45.1%) have no classes. (b) The super classes of all remaining 263 metabolites.

## Notes

https://jaspershen.github.io/metID/index.html

